# SARS-CoV-2 Beta Variant Substitutions Alter Spike Glycoprotein Receptor Binding Domain Structure and Stability

**DOI:** 10.1101/2021.05.11.443443

**Authors:** Daniel L. Moss, Jay Rappaport

## Abstract

The emergence of severe acute respiratory syndrome-related coronavirus 2 (SARS-CoV-2) and the subsequent COVID-19 pandemic has visited a terrible cost on the world in the forms of disease, death, and economic turmoil. The rapid development and deployment of extremely effective vaccines against SARS-CoV-2 have seemingly brought within reach the end of the pandemic. However, the virus has acquired mutations; and emerging variants of concern (VOC) are more infectious and reduce the efficacy of existing vaccines. While promising efforts to combat these variants are underway, the evolutionary pressures leading to these variants are poorly understood. To that end, here we have studied the effects on the structure and function of the SARS-CoV-2 spike glycoprotein receptor-binding domain of three amino-acid substitutions found in several variants of concern, including alpha (B.1.1.7), beta (B.1.351), and gamma (P.1). We found that these substitutions alter the RBD structure, stability, and ability to bind to ACE2, in such a way as to possibly have opposing and compensatory effects. These findings provide new insights into how these VOC may have been selected for infectivity while maintaining the structure and stability of the receptor binding domain.

## Introduction

The emergence of SARS-CoV-2 in late 2019 and its subsequent spread around the world has caused the deadliest airborne pandemic in the United States, recently surpassing the 1918 influenza pandemic nearly a century ago(1). The international scientific community has risen to the challenge of combating SARS-CoV-2 and COVID-19. The year 2020 ended with the fastest development of vaccine candidates, starting with the genetic sequence of the virus being reported(2) to human trials of novel mRNA-based vaccines within three months. Now there are three SARS-CoV-2 vaccines approved for use within the United States and many more next-generation and pan-coronavirus vaccines currently in development. These advances have made substantial contributions to the control of the COVID-19 pandemic within the United States. Despite multiple manufacturers receiving emergency use authorization, and an unprecedented vaccination campaign, significant challenges remain including uncertainty regarding durability, vaccination hesitancy, limited access to healthcare among disadvantaged persons, as well as the continued emergence of VOC. Our ultimate success in quelling this pandemic may ultimately lie in our ability, not only to characterize new variants but to be able to predict the emergence of new variants. Such advances will require an increased understanding of evolutionary pressures and constraints on viral variation.

Three SARS-CoV-2 lineages, the alpha variant lineage B.1.1.7 (or 501Y.V1) first identified within the United Kingdom, the beta variant lineage B.1.351 (or 501Y.V2) identified in South Africa, and the gamma variant lineage P.1 (or 501Y.V3) identified in Brazil, have been demonstrated to possess increased infectivity(3) and in the case beta and gamma exhibit reduced neutralization by antibodies reacting with the cognate regions of the spike protein within the original Wuhan strain of SARS-CoV-2(4–6). The alpha variant possesses the N501Y substitution within the spike glycoprotein receptor binding domain (RBD) which has been shown to enhance binding to ACE2, the entry receptor for SARS-CoV-2(7–9). The beta and gamma variants possess N501Y as well as substitutions at two other sites within the RBD, E484K, and K417N in beta and K417T in gamma(10). These RBD substitutions present in the spike protein of the B.1.351 and P.1 variants have been shown to reduce the binding and neutralization of mRNA vaccine-induced antibodies as well as potent human monoclonal antibodies(11).

The consequences of the K417N, E484K, and N501Y substitutions on RBD-ACE2 interactions have also been examined, with the increased infectivity of the alpha variant resulting from enhanced binding to ACE2 when the RBD N501Y substitution is present(9). The E484K substitution has been shown to enhance ACE2 binding(12) and reduce the efficacy of neutralizing antibodies(13). A recent study examined the effects of the K417N substitution on ACE2 binding and antibody interactions using molecular dynamics and found that K417N disrupts RBD-ACE2 interactions, as well as interactions with a monoclonal antibody(14). However, the effects of these substitutions on the structure of the RBD itself have not been examined. Based on the nature of these substitutions, including residue changes in charge or polar to non-polar substitutions, we hypothesized that the K417N, E484K, and N501Y substitutions alter RBD structure and stability as well as ACE2 binding interactions. We studied those changes in single-substitution RBD variants as well as in the RBD containing all three substitutions using molecular dynamics and biophysical approaches. Our data suggest that these VOC substitutions significantly alter RBD structure and stability, with consequences for ACE2 binding and proteolytic susceptibility, having potentially opposing consequences for the fitness of new variants. These findings have implications for viral evolution as well as the design of subunit vaccine candidates.

## Results

### RBD Variant of Concern Substitutions Alter RBD Structure *in silico*

We began our studies on RBD variant of concern substitutions with molecular dynamics to investigate whether single substitutions or all three substitutions found within the beta variant would alter RBD structure. We used homology modeling with residues 319 – 541 of the trimeric spike glycoprotein as a template (PDB 6VXX) to generate structures of the RBD(15, 16). The resulting model of the wild-type/Wuhan strain RBD was used as a template for further modeling of RBD variants containing single substitutions as well as all three substitutions (K417N/E484K/N501Y) found within the beta variant. Molecular dynamic simulations were performed with the GROMACS 2020.5 package (17) using the CHARMM3636 or ff14SB force fields(18, 19). After solvation and neutralization, systems were minimized and equilibrated before undergoing a 25-nanosecond production run within the NPT ensemble. From these trajectories, we observed no large differences in root mean square deviation (RMSD) relative to the starting structure for any of the RBD variants (Figure 1A and 1B). In the CHARMM36 force field, the RMSD for the N501Y and K417N/E484K/N501Y RBD variants began to diverge relative to the other RBD variants after about 15 ns. The RMSD for the K417N/E484K/N501Y variant quickly converged with the others by the end of the simulation run whereas in the ff14SB force field the RMSD was consistent across all RBD variants. Changes in residue-specific fluctuations were observed for K417N, N501Y, and K417N/E484K/N501Y RBD variants in the CHARMM36 force field (Figure 1C) but were absent in the ff14DB forcefield except for K417N/E484K/N501Y exhibiting a slight increase in overall residue fluctuations (Figure 1D). We also observed a decrease in residue RMSF values around the region of residues 468 – 488 in the ff14SB forcefield for the E484K, N501Y, and K417N/E484K/N501Y variants that were not seen for the wild-type and K417N RBD. Overall, in the CHARMM36 force field we observed VOC substitutions causing more variation in RMSD and RMSF than in the ff14SB force field.

**Figure 1:**
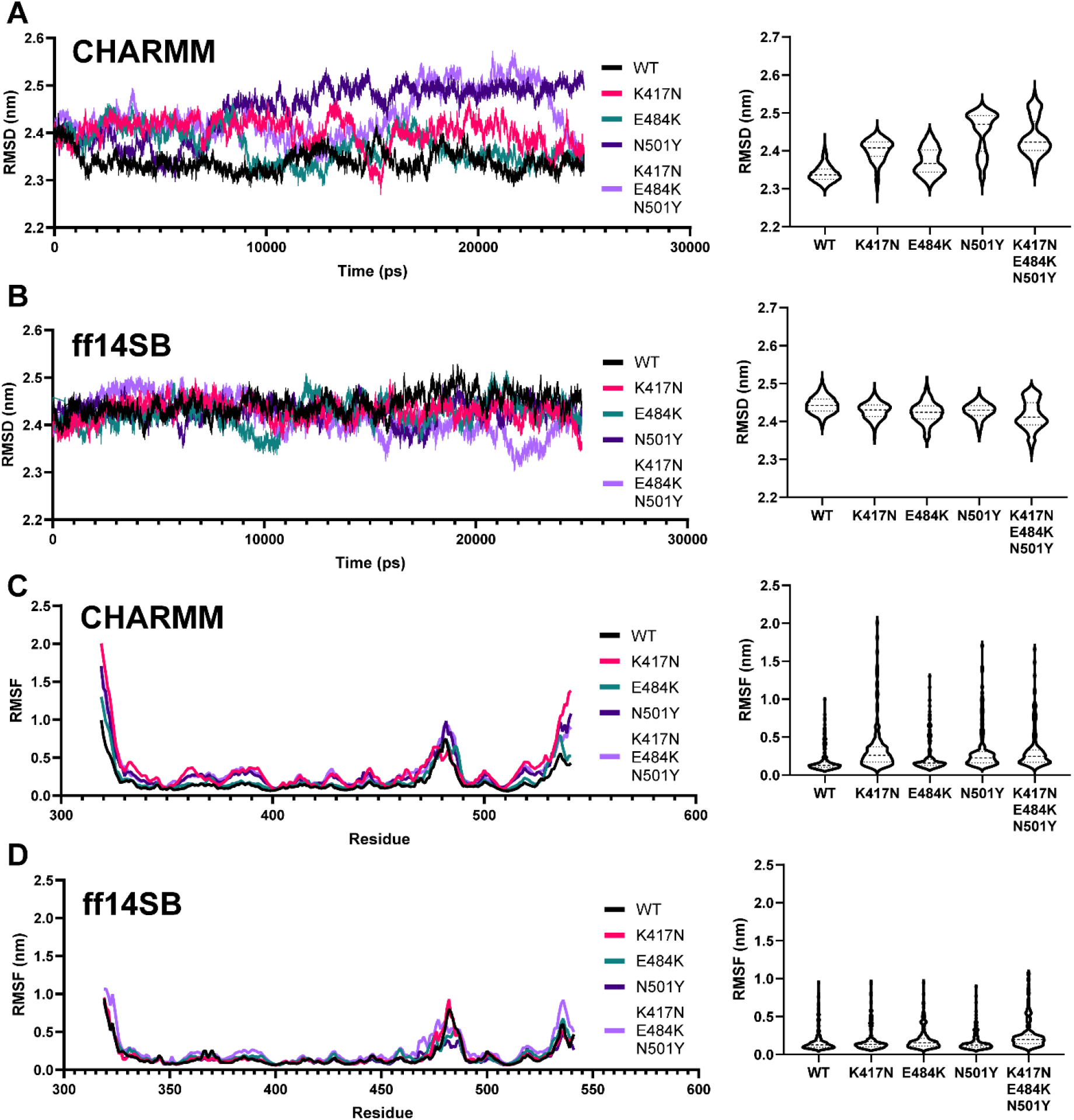
Molecular Dynamic Simulations of SARS-CoV-2 Receptor Binding Domain Suggests that Variant-of-Concern Substitutions Alter RBD Structure. A & B. Root mean square deviation plotted as a function of time (left) and averaged as violin plot (right) for each RBD variant simulated in the CHARMM36 (A) or ff14SB (B) force fields. C & D. Root mean square fluctuation plotted as a function of time (left) or averaged and shown as a violin plot (right) in the CHARMM36 (C) or ff14SB (B) force fields.

Next, we examined hydrogen bonding and radius of gyration for all RBD variants in both force fields. Hydrogen bond content was lower in the CHARMM36 forcefield (Figure 2A) compared to the ff14SB force field (Figure 2B) while in both force fields the K417N substitution increased average hydrogen bond content relative to all other RBD variants except for E484K in the ff14SB force field. In the CHARMM36 force field, all RBD variants exhibited an increase in the radius of gyration, a measure of the compactness of the molecule, relative to the wild-type RBD over throughout the production run with the greatest increases observed for the N501Y and K417N/E484K/N501Y RBD variants (Figure 2C). In the ff14SB force field, the radius of gyration was overall consistent for all RBD variants throughout the production run with some slight increases observed for the K417N and K417N/E484K/N501Y RBD variants within the initial 10 ns of the simulation (Figure 2D). Taken together these observations suggest that the CHARMM36 force field predicts that VOC substitutions will have a more disruptive effect on the RBD structure compared to the ff14SB force field.

**Figure 2:**
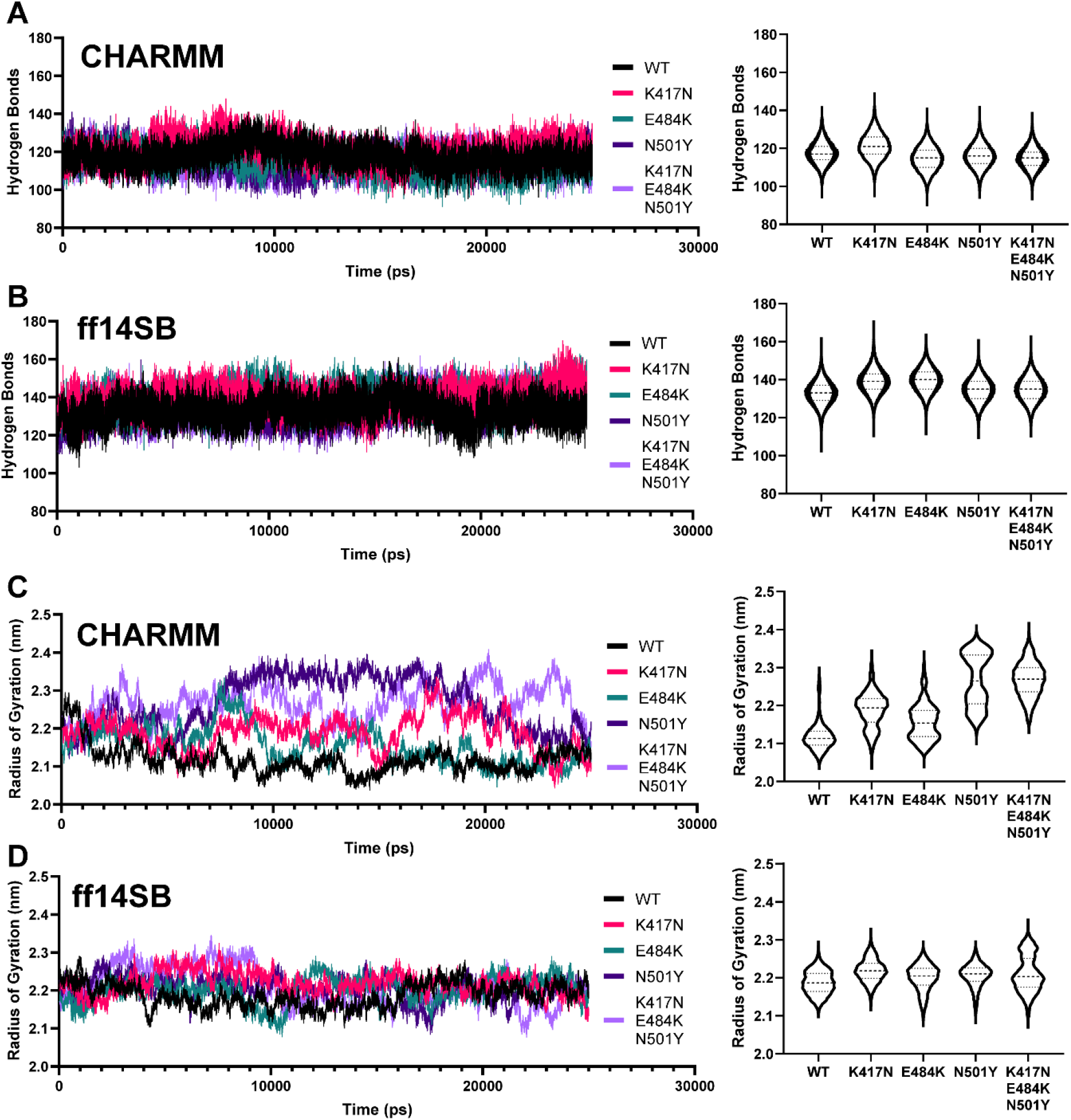
Molecular Dynamic Simulations of SARS-CoV-2 Receptor Binding Domain Suggests that Variant-of-Concern Substitutions Alter RBD Structure. A & B. Total number of hydrogen bonds within the RBD for each variant simulated plotted as a function of time (left) and averaged and plotted in a violin plot (right) in the CHARMM36 (A) or ff14SB (B) force fields. C & D. Radius of gyration plotted as a function of time (left) and averaged as violin plot (right) for each RBD variant in the CHARMM36 (C) or ff14SB (D) force fields.

To identify common structural changes and differences in residue contacts we extracted frames from the simulation trajectories and examined the RBD structure near the sites of the beta variant substitutions at the beginning (100 ps, green) and end (25000 ps, cyan) of the production run in both force fields. In the CHARMM36 force field for the wild-type RBD, we observe K417 and E484 interacting with solvent while N501 forms a hydrogen bond with the polypeptide backbone as well as a hydrogen bond between R403 and E406 (Figure 3A). At the end of the simulation, we observed changes in secondary structure and repositioning of the 468 – 488 loop. When K417 is substituted for asparagine, we observed N417 interacting with E406 at the beginning of the simulation but by the end, N417 has formed solvent interactions while a significant rearrangement of the 468 – 488 loop is observed with E484 forming a hydrogen bond with R403 (Figure 3B). This change in conformation occurs within the first 5 ns of the simulation as shown by the rapid decrease in distance between residues 403 and 484 (Figure 3G) and residues 417 and 484 (Figure 3H). For the E484K variant, both the early and late structures are very similar with some changes in the 468 – 488 loop secondary structure (Figure 3C). For the N501Y variant, we observed that the substitution at position 501 eliminated the backbone interactions observed for N501 in the wild-type structure as well as changes in the position of the 468 – 488 loop (Figure 3D). We also observed changes in the positioning of K417, E406, and R403 while the hydrogen bond between R403 and E406 was maintained. For the K417N/E484K/N501Y RBD variant, we observed the same R403, E406 interactions we observed in the other RBD variants (Figure 3E). However, N417 maintained an interaction with E406 that was not observed in the K417N trajectory. By the end of the simulation for K417N/E484K/N501Y we observed the 468 – 488 loop backbone carboxyl groups interacting with arginine 457 (Figure 3F), a shift in conformation that occurs after about 14 ns in the simulation trajectory and can be seen by changes in the distances between residues 403 and 484 (Figure 3G) as well as residues 417 and 484 (Figure 3H). Overall, the observed changes in residue contacts for the K417N and K417N/E484K/N501Y RBD variants reduced the RMSD, relative to the starting structure, of the 468 – 488 loop compare to the wild-type, E484K and N501Y variants (Figure 3I). In the CHARMM36 force field we observed stabilization of the 468 – 488 loop in both the K417N and K417N/E484K/N501Y variants by hydrogen bonding between R403 for K417N and R457 for K417N/E484K/N501Y.

**Figure 3:**
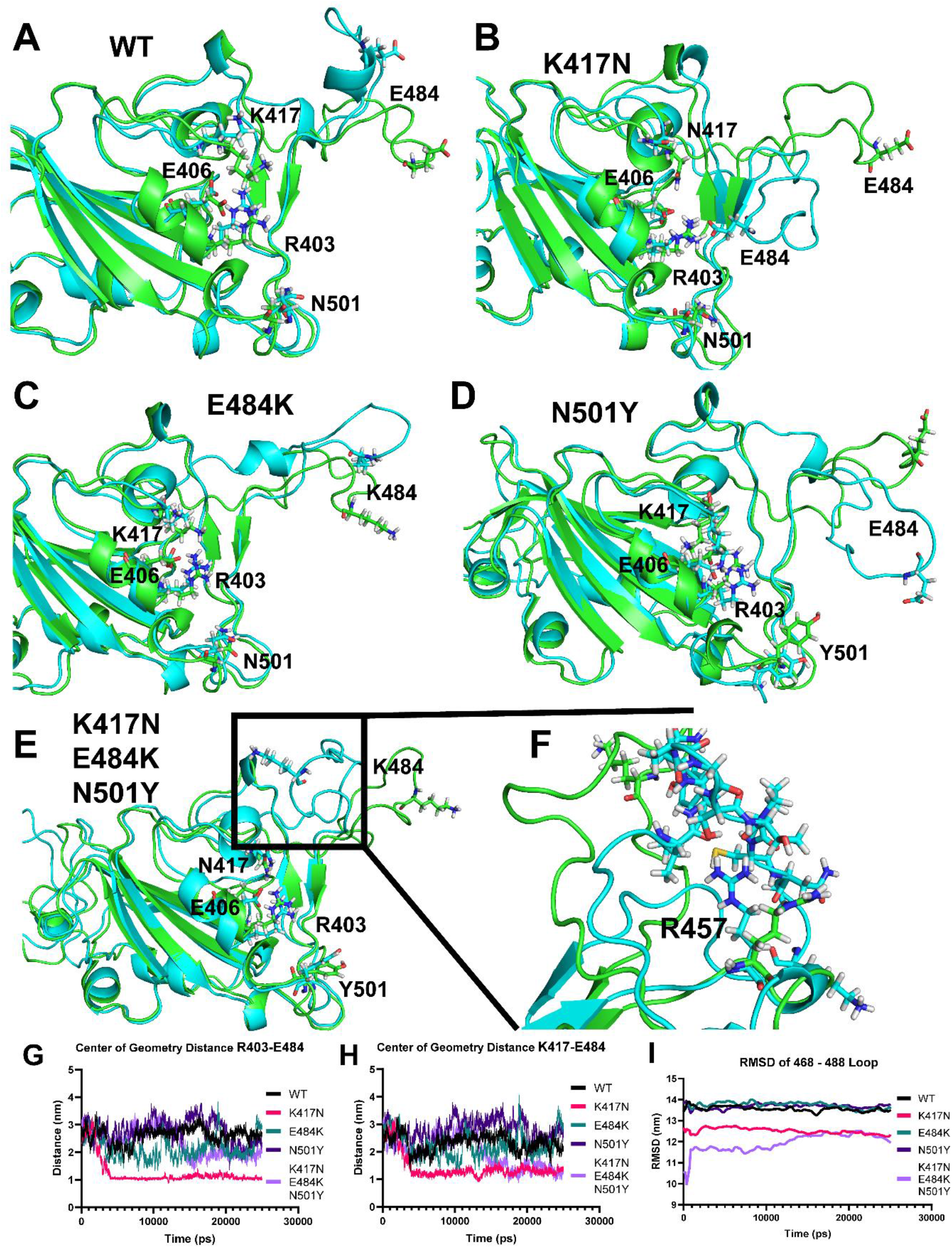
CHARMM36 specific changes in RBD residue contacts. A - E. RBD coordinates extracted from the molecular dynamics trajectories of the wild-type (A), K417N (B), E484K (C), N501Y (D) and K417N/E484K/N501Y (E) RBD variants in the CHARMM36 force field. The substituted residues as well as important hydrogen bonding pairs are shown. F. Closer view of the 468 – 488 region of the K417N/E484K/N501Y RBD shown in E to highlight the interaction between the side chain of arginine 457 and the backbone carboxyl groups of the 468 – 488 loop. G. Center of geometry distance in nm between residue R403 and E484 plotted as a function of time for all RBD variants in the CHARMM36 force field. H. Center of geometry distance in nm between residue K417 and E484 plotted as a function of time for all RBD variants in the CHARMM36 force field. I. RMSD of the 468 – 488 loop relative to the starting structure for all RBD variants plotted as a function of time.

We next compared structures extracted from trajectories in the ff14SB force field. For the wild-type RBD, we observed K417 forming a hydrogen bond with E406 at both time points (Figure S1A). Here we also observed a transition in the 468 – 488 loop from a more solvent-exposed position to one forming primarily backbone contacts rather than a hydrogen bond between E484 and R403 which remain relatively far apart throughout the simulation run (Figure S1F). However, this conformation does not appear to be occupied for a long period of time as the RMSD for this region is elevated compared to K417N and K417N/E484K/N501Y (Figure S1H). We do observe N417 participating in electrostatic interactions with R403 and E406 (Figure S1A). For the K417N variant in ff14SB, we do not observe a similar conformational change to that observed in CHARMM36 but rather we observed the formation of a hydrogen bond between E484 and R457 and this region exhibits a reduced RMSD relative to the other variants except K417N/E484K/N501Y (Figure S1B and S1H). Changes in the structure of the E484K variant in ff14SB are similar to those observed for the wild-type RBD with the exception of the 468 – 488 loop which does not form the same backbone contacts seen for the wild-type RBD (Figure S1C). For the N501Y variant in ff14SB, we do not observe the same K417 – E406 interaction seen in the wild-type RBD (Figure S1D). For the K417N/E484K/N501Y variant in ff14SB, we observed changes in secondary structure within the 468 – 488 loop that may explain the reduced RMSD in this region relative to the other variants except K417N (Figure S1E). We also observed electrostatic interactions between N417, R403, and E406. As in CHARMM36, in the ff14SB force field we observed stabilization of the 468 – 488 loop for the K417N and K417N/E484K/N501Y RBD variants by different mechanisms. For K417N, a hydrogen bond between R457 and E484 while for K417N/E484K/N501Y it appears that helical formation in the 468 – 488 loop is responsible for its relative stability compared to the wild-type RBD.

### RBD Variant of Concern Substitutions Alter RBD Stability and ACE2 Binding Affinity

Molecular dynamic simulations of RBD variants suggested that variant of concern substitutions alter RBD structure and hydrogen bonding. Therefore, we hypothesized that these RBD substitutions alter RBD stability and resistance to unfolding. To test this we measured guanidine-induced unfolding of RBD variants using a fluorescence-based unfolding assay (20, 21). Bis-ANS fluorescence was measured for RBD variants in the presence of increasing guanidine-HCl concentration (Figure 4A). Fluorescence data were normalized and unfolding curves were fit by non-linear regression to estimate the free energy of unfolding (Figure 4B)(22). We observed that the E484K substitution significantly destabilized the RBD while the N501Y and K417N substitutions significantly stabilized the RBD relative to wild-type (Figure C). We observed no difference in unfolding energy when all three beta variant substitutions were present in the RBD compared to the wild-type protein. Changes in m-values for RBD variants corresponded with changes in folding free energy. We observed a decrease in the m-value for the E484K variant suggesting a reduction in the difference between solvent accessible surface area between the folded and unfolded states compared to the wild-type RBD (Figure 4D)(23). For both the K417N and N501Y variants we observed increased m-values suggesting that the unfolded and folded states exhibit greater differences in exposed surface area upon unfolding relative to the wild-type RBD. A summary of the parameters fit by non-linear regression are shown in Table 1. Changes in resistance to denaturant-induced unfolding did not correspond with protein melting temperatures as measured by a fluorescence melting assay that we have applied to the study of ovalbumin variants (Figure 4E)(20). We observed a reduction in the melting temperature of the beta variant RBD (Figure 4F).

**Figure 4:**
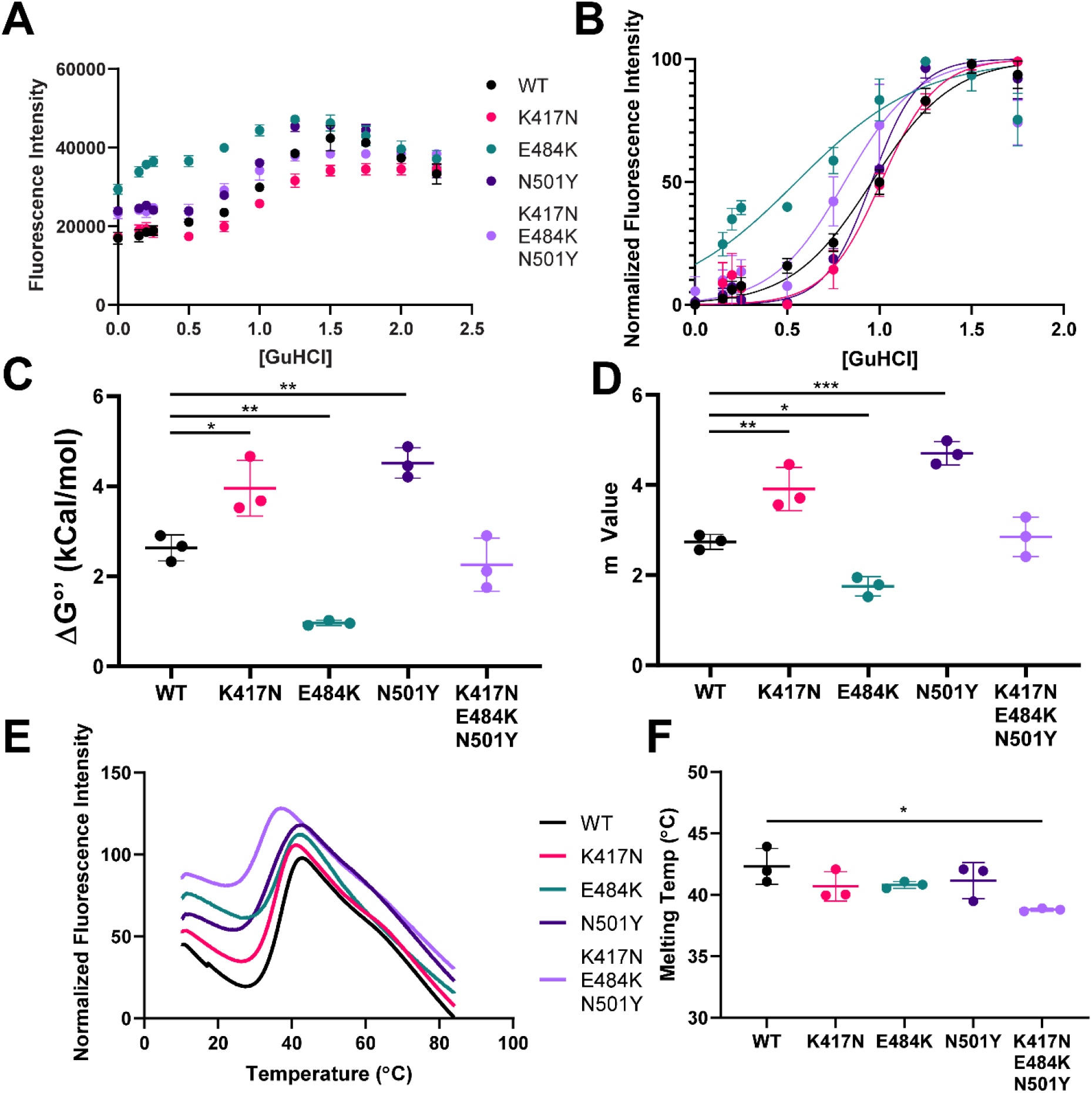
RBD Variant of Concern Substitutions Alter RBD Stability. A. Raw Bis-ANS fluorescence data as a function of guanidine-HCl concentration for the indicated RBD variant tested. B. Unfolding curves for the indicated RBD variant generated by monitoring Bis-ANS fluorescence as a function of guanidine-HCl concentration. Non-linear regression was used to analyze unfolding data. Data points indicated the mean measurement of three replicates while error bars indicate standard deviation. C. Calculated ΔG’° values from non-linear regression analysis for each RBD variant were compared by one-way ANOVA and Tukey’s test for multiple comparisons. P value less than 0.05 is indicated by *, p value less than 0.01 is indicated by **. Error bars indicate standard deviation. D. Non-linear regression m values for each RBD variant were compared by one-way ANOVA and Tukey’s test for multiple comparisons. P value less than 0.05 is indicated by *, p value less than 0.01 is indicated by ** and p value less than 0.001 is indicated by ***. Error bars indicate standard deviation. Guanidine unfolding data are representative of three technical replicates using the same preparation of RBD. E. Temperature induced unfolding of RBD variants. Bis-ANS fluorescence data as a function of temperature is plotted for the indicated RBD variant. The peak of each curve was taken as the melting temp. F. Melting temperatures for the indicated RBD variant were compared by one-way ANOVA and Tukey’s test for multiple comparisons. P value less than 0.05 is indicated by *. Error bars indicate standard deviation. Each RBD variant was measured in triplicate using a unique preparation of RBD.

**Table 1.** Summary of parameters fit by non-linear regression.

| **RBD Variant** | **$\Delta G^{\circ}$ (SEM)**<br>**(kCal/mol)** | **m-value (SEM)** | **Mid-point [GuHCl] (SEM)** |
| --- | --- | --- | --- |
| **WT** | 2.610 (± 0.173) | 2.718 (± 0.175) | 0.960 (± 0.017) |
| **K417N** | 3.962 (± 0.463) | 3.922 (± 0.452) | 1.010 (± 0.020) |
| **E484K** | 0.966 (± 0.132) | 1.752 (± 0.221) | 0.551 (± 0.045) |
| **N501Y** | 4.516 (± 0.443) | 4.715 (± 0.455) | 0.958 (± 0.013) |
| **K417N/E484K/N501Y** | 2.452 (± 0.409) | 3.018 (± 0.490) | 0.812 (± 0.039) |

As discussed above, it has been reported that the substitutions to the spike protein observed in the alpha and beta variants exhibit a higher binding affinity for ACE2. We sought to replicate these observations and examine the effects of the individual K417N and E484K substitutions on ACE2 binding affinity. It has also been reported that K417N reduces RBD binding affinity for ACE2 by both computational and experimental investigation(14, 24). To test the effects of RBD substitutions on ACE2 binding we utilized a binding competition assay where RBD variants compete with horseradish peroxidase-labeled wild-type RBD (HRP-RBD) for binding to ACE2 adsorbed to a microplate. As RBD variants are diluted HRP-RBD can outcompete for binding to ACE2 resulting in increased absorbance at 450 nm after plate development (Figure 5A). These data were analyzed by non-linear regression to calculate LogIC50 values. We found that the K417N substitution significantly reduced ACE2 binding as less dilution was needed for HRP-RBD to outcompete for ACE2 binding (Figure 5B). E484K exhibited a slightly higher affinity for ACE2 when compared to the wild-type RBD but this did not reach the level of statistical significance. The N501Y substitution resulted in increased ACE2 affinity consistent with previous reports and the presence of all three variant of concern substitutions exhibited the greatest increase in ACE2 affinity compared to wild-type RBD (Figure 5B). These results were consistent with previously reported observations of the effects of variant of concern substitutions on RBD-ACE2 binding(8, 9, 14, 24).

**Figure 5:**
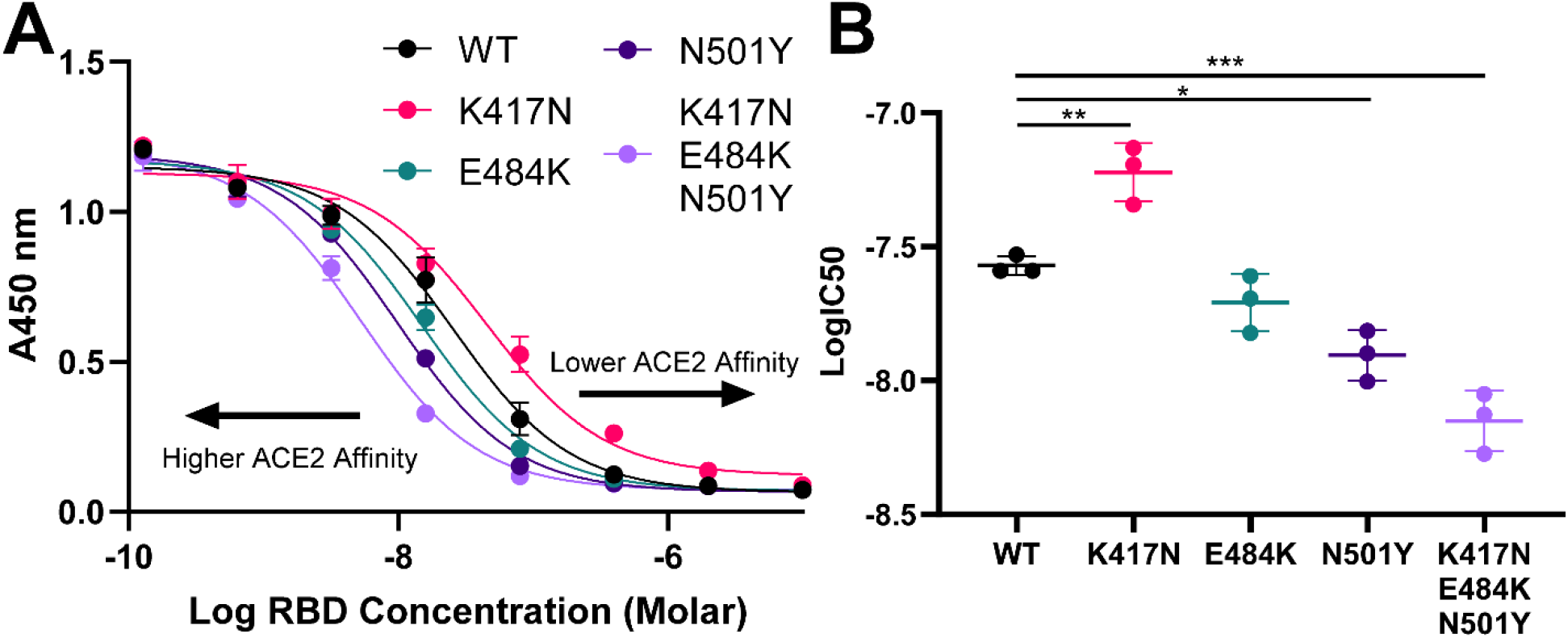
RBD Variant of Concern Substitutions Alter ACE2 Binding Affinity. A. Inhibition curves for each indicated RBD variant. As the RBD variant is diluted out HRP labeled wild-type RBD can outcompete for binding to ACE2 resulting in an increase in absorbance at 450 nm. Data were analyzed by non-linear regression. B. LogIC50 values from A were compared by one-way ANOVA and Tukey’s test for multiple comparisons with asterisks indicating p value: * for < 0.05, ** for < 0.01, *** for < 0.001. Data shown are representative of three independent experiments using the same preparation of RBD. Error bars indicate standard deviation.

### RBD Variant of Concern Substitutions Alter RBD Proteolytic Susceptibility

Molecular dynamic simulations and results from unfolding studies presented here lead us to the conclusion that RBD variant of concern substitutions alter RBD structure and stability. Changes in protein structure and stability are associated with changes in proteolytic resistance(20, 21, 25, 26). Based on these previous observations and those we have reported here thus far, we hypothesized that RBD substitutions alter proteolytic susceptibility in accordance with changes in stability. Substitutions that stabilize the RBD increase proteolytic resistance, while destabilizing substitutions will decrease proteolytic resistance. To test this, we performed limited proteolysis experiments using the lysosomal protease cathepsin S at pH 5.6. Proteolysis reactions were sampled after 0, 15, 30, and 60 minutes of incubation and analyzed by SDS-PAGE and Coomassie staining (Figure 6A). Gels were imaged and intensities of the RBD bands were measured with gel analysis software. Band intensities were normalized to the 0 minutes time point and compared by two-way ANOVA. We observed that the K417N substitution significantly increased proteolytic resistance at all time points compared to the wild-type RBD (Figure 6B). The E484K substitution significantly decreased proteolytic resistance compared to the wild-type RBD except after 60 minutes of incubation. The N501Y substitution exhibited increased resistance to proteolysis by cathepsin S only after 60 minutes of incubation relative to the wild-type RBD. The presence of all three substitutions resulted in a large increase in cathepsin S proteolytic susceptibility after 15 and 30 minutes but degradation was similar to that seen for wild-type RBD after 60 minutes (Figure 6B). Taken together these results demonstrate that RBD variant of concern substitutions significantly alter RBD proteolytic susceptibility.

**Figure 6:**
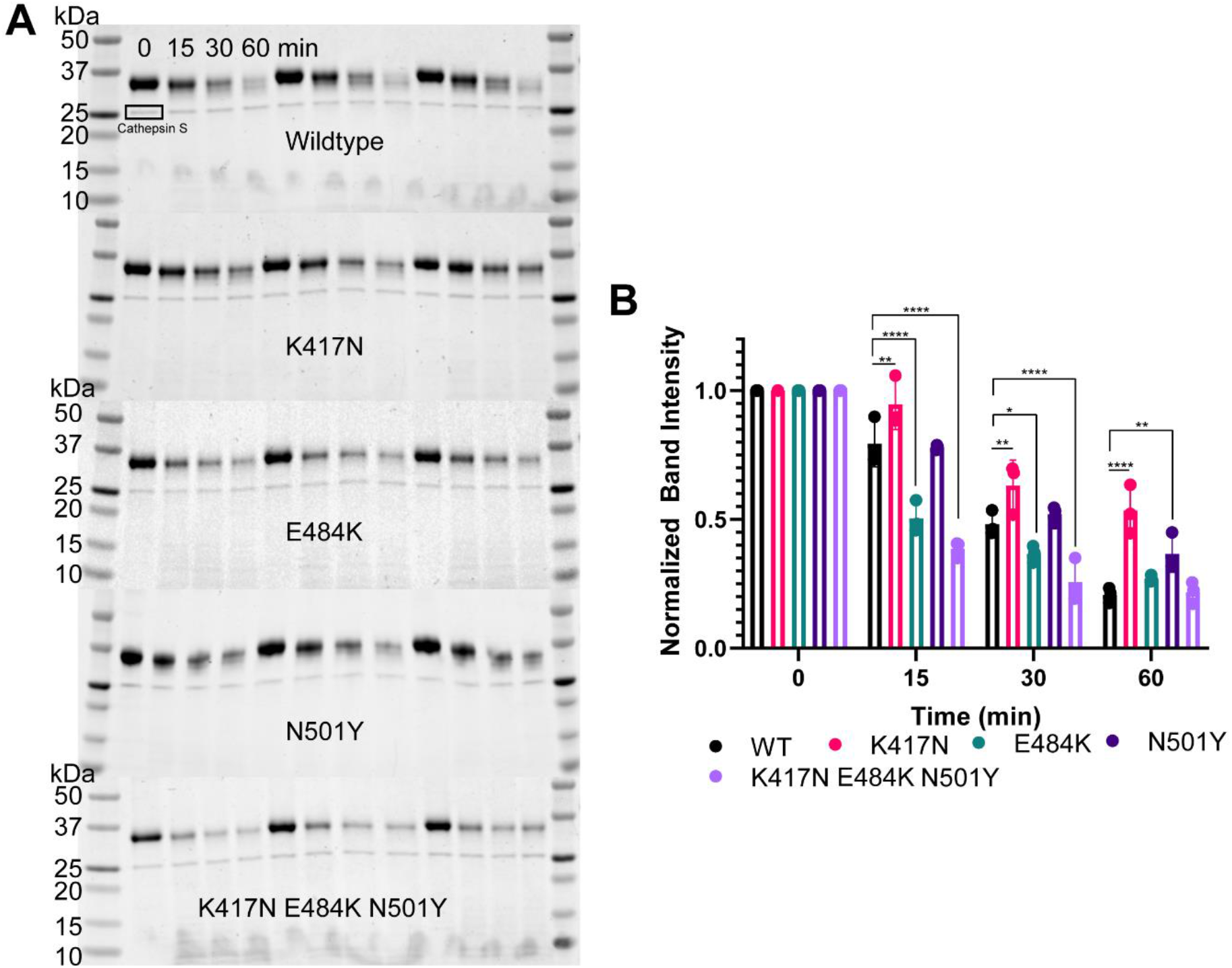
RBD Variant of Concern Substitutions Alter Susceptibility to Limited Proteolysis by Cathepsin S. A. SDS-PAGE and Coomassie staining of limited proteolysis reactions for each RBD variant indicated run in triplicate using the same preparation of RBD. B. Band intensities were calculated from A and normalized to the 0 min time point. Mean intensities for each replicate are shown, error bars indicate standard deviation and n = 3. Mean values were compared by two-way ANOVA and Dunnett’s test for multiple comparisons asterisks indicating p value: * for < 0.05, ** for < 0.01 and **** for < 0.0001.

## Discussion

In this study, we tested the hypothesis that SARS-CoV-2 beta variant substitutions within the spike glycoprotein RBD alter the RBD structure, stability, and ACE2 binding affinity. We studied the RBD by molecular dynamics simulation and found that the K417N substitution alone as well as the presence of all three substitutions, K417N/E484K/N501Y, within the beta variant RBD may alter the flexibility of a loop region spanning residues 468 – 488 that forms contacts with ACE2. A similar molecular dynamic simulation analysis of this region was reported for the wild-type RBD and the E484K, N501Y, and the K417N/E484K/N501Y substitutions, albeit with different observations to those reported here(27). The authors observed that the E484K and N501Y substitutions alone increased flexibility relative to wild-type while both E484K and N501Y together along with all three substitutions (K417N/E484K/N501Y) did not alter flexibility compared to the wild-type RBD. Our results differ here; in the CHARMM36 force field, we observed minor changes in the RMSF values for the 468 – 488 region for all RBD variants while in the ff14SB force field the K417N substitution did not result in large RMSF changes while the E484K, N501Y and K417N/E484K/N501Y substitutions reduced RMSF values relative to the wild-type RBD which, except for the K417N/E484K/N501Y RBD variant, were not consistent with the results from the previous study. While the previous study performed their simulations in the ff14SB forcefield, differences in the equilibration parameters and production run time may account for these discrepancies. It is challenging to say which results are more plausible and comparison to experimental data is difficult as there are no available structures of the RBD in the unbound state.

The major differences observed here between CHARMM36 and ff14SB are the distances between residues 403 and 484 as well as 417 and 484. In CHARMM36 for K417N, and less so for K417N/E484K/N501Y, these residues quickly form close contacts that are not observed for the other variants while in ff14SB these residues are too far apart to participate in meaningful structural interactions. Interestingly in both CHARMM36 and ff14SB the 468 – 488 loop exhibits reduced RMSD for the K417N and K417N/E484K/N501Y variants relative to the other three (wild-type, E484K and N501Y). Upon examination of the coordinates within the respective trajectories, it appears this observation arose through different structural mechanisms. For K417N in the CHARMM36 force field, the hydrogen bond between R403 and E484 appears to be limiting the flexibility of the 468 – 488 loop while in ff14SB the R403 – E484 hydrogen bond is not observed and the 468 – 488 loop appears to be stabilized by a hydrogen bond between R457 and E484. A similar interaction between R457 and the backbone of the 468 – 488 loop stabilizes this region in the K417N/E484K/N501Y variant in the CHARMM36 force field. Others have recently reported that the CHARMM36 force field favors more disordered protein conformations while the ff14SB force field favors the native state and a stabilization of alpha helical structures which may explain the helix formation within the 468 – 488 loop of the K417N/E484K/N501Y in ff14SB(28). The biases reported for the CHARMM36 and ff14SB force fields may also explain our observations of the differences in RMSD and radius of gyration we observed between these two force fields. CHARMM36 predicted greater changes in RMSD and radius of gyration resulting from VOC substitutions than ff14SB.

In a binding competition assay, the N501Y and K417N/E484K/N501Y variants all exhibited a higher binding affinity for ACE2 while the K417N substitution alone exhibited reduced binding affinity. The E484K substitution caused a modest but insignificant increase in affinity as measured by the competition assay employed here and other studies have reported the E484K substitution results in a negligible increase in affinity(29). Nonetheless, our observations are all in agreement with numerous previous reports on the effects of these substitutions on RBD-ACE2 binding(9, 12, 14, 24, 30–33). Reduced binding to ACE2 in K417N may result from a decrease in the flexibility of the 468 – 488 region that undergoes a conformational change when bound to ACE2 in the RBD variant containing only the K417N substitution. Analysis of the simulation trajectories in the CHARMM36 force field suggests that glutamate 484 forms a hydrogen bond interaction with arginine 403 when only the K417N substitution is present as these residues are much further apart in simulation data generated from the wild-type and E484K RBD variant. The presence of the hydrogen bond between E484 and R403 results in a decrease in the RMSD of the loop spanning residues 468 – 488 in K417N when compared to the wild-type and E484K RBD variants. In the ff14SB force field, the 468 – 488 loop in the K417N appears to be stabilized by a hydrogen bond between R457 and E484. The 468 – 488 loop region has been reported to undergo a structural transition that is important for ACE2 binding(34). Therefore, a hydrogen bond between E484 and R403, or E484 and R457, that restricts the flexibility of the 468 – 488 loop may be responsible for the reduction in ACE2 binding affinity we observed here but more experiments would be necessary to determine this. More extensive simulation experiments in a previous study have shown that the K417N substitution may abolish a salt bridge of K417 with an aspartate residue at position 30 within ACE2 and therefore reduce the strength of binding interactions(14). If this alone is enough to explain the observed reduction in ACE2 binding by the K417N variant or if it simply contributes remains to be determined.

The utility of the K417N substitution to the virus remains unclear. It has not been reported alone in other variants and this may be due to its negative effects on ACE2 binding reported here and by others(14). K417N also appears to reduce the strength of antibody-RBD interactions and may have been selected for immune evasion despite its potential negative effects on transmissibility. Our data suggest that K417N alone alters the RBD structure in such a way that conformational changes necessary for ACE2 binding may be disfavored relative to the wild-type or other variants. We also observed that the E484K substitution alone significantly destabilizes the RBD by both denaturant-induced unfolding and limited proteolysis. Therefore, the stabilizing effects of the K417N substitution may be necessary to offset the negative structural effects of the E484K substitution. The N501Y substitution also stabilized the RBD in our studies and the iota variant (B.1.526) has been identified in New York that possesses both the N501Y and E484K substitutions(35). The K417N and E484K substitutions have not yet been reported to exist alone in a SARS-CoV-2 variant and it is possible their beneficial effects for immune evasion in the case of both K417N and E484K or ACE2 binding in the case of E484K are outweighed by negative structural effects. However, the caveat here is that the RBD does not exist in isolation but rather as part of the much larger spike glycoprotein, but the effects of these substitutions on structure and stability may still be relevant for understanding the evolution of SARS-CoV-2.

The presence of K417N alone and N501Y alone significantly stabilized the RBD relative to wild-type and it is unclear from the simulation data how this was achieved for the N501Y substitution found in the alpha variant. Surprisingly the E484K substitution alone significantly destabilized the RBD relative to wild-type. Our simulation data suggest that E484K alone reduces overall hydrogen bond content in the RBD. Reduced hydrogen bond content may explain the observed decrease in stability and suggests that the elimination of specific interactions between residue 484 and other RBD residues alone cannot explain the observed reduction in RBD stability. There are, however, limitations to the bis-ANS unfolding assay used in this study. This assay is based on the measurement of bis-ANS fluorescence as a function of denaturant-induced protein unfolding. The fluorescent dye bis-ANS is quenched by water and binds to hydrophobic regions of proteins increasing fluorescence. Due to the binding nature of bis-ANS, the dye may unbind before the protein is completely unfolded as has been seen for the unfolding of ovalbumin(20) and was observed here. Fluorescence signal peaked at 1.25 to 1.5 M guanidine HCl and began to decrease at the RBD became more solvent-exposed. As denaturant concentration increases eventually hydrophobic regions of proteins will become completely exposed to water solvent and the dye will unbind and be quenched. However, as was seen for ovalbumin and was reported in this study bis-ANS binding in the presence of denaturant can still be informative of protein stability. It is also possible that amino acid substitutions may alter the binding affinity of the dye rather than protein stability. We do not consider that to be a major factor for our results as the substitutions studied here were of polar residues or, in the case of N501Y, surface-exposed and therefore we would not expect those substitutions to alter dye-binding sites. Furthermore, the results from the bis-ANS unfolding experiments were corroborated by limited proteolysis with cathepsin S. The stabilized K417N and N501Y variants were more resistant to cathepsin S proteolysis than the destabilized E484K variant.

In summary, we have reported here that RBD variant of concern substitutions alter RBD structure and stability with K417N and N501Y increasing stability while E484K reduces stability. All three substitutions together, as found in the beta variant, exhibit similar stability to the wild-type RBD albeit with a possibly more open conformation and significantly higher ACE2 binding affinity that is greater than the sum of its parts. Taken together our findings support the notion that the evolution of the SARS-CoV-2 RBD has been guided by pressure for increased ACE2 binding and immune evasion within the constraints of maintaining RBD structure for optimal interactions with the ACE2 receptor.

## Materials and Methods

### Molecular Dynamics Simulations

Molecular dynamic simulations were performed with the GROMACS 2020.5 package(17, 36) with the CHARMM3636 all-atom force field(18) or the ff14SB force field(19), and the TIP3P water model(37). The wild-type SARS-CoV-2 spike glycoprotein receptor binding domain (residues 319 – 541) and variants were modeled from the full-length spike glycoprotein Cryo-EM structure PDB entry 6VXX(16) using SWISS-MODEL(15). RBD models were solvated in a dodecahedral box with a minimum protein to edge distance of 1.5 nm and 37162 water molecules. The system charge was neutralized with approximately 105 Na^+^ and 113 Cl^−^ ions at a concentration of 0.15 M. Assembled systems were minimized for 10 ps and then equilibrated for 100 ps in the *NVT* ensemble followed by further equilibration for 1000 ps with harmonic position restraints on heavy protein atoms (1000 kJ mol^−1^ nm^−2^) and Berendsen coupling(38) to maintain temperature and pressure (P ~ 1 bar, and T ~ 300 K). Systems were further equilibrated for 25 ns in the *NPT* ensemble followed by three individual 10 ns production runs for each variant also in the *NPT* ensemble. Trajectory data were saved every 1 ps for CHARMM36 simulations and every 0.5 ps for ff14SB simulations. Analysis was performed with GROMACS, VMD 1.9.3(39), PyMOL, and GraphPad Prism.

### Protein Expression and Purification

The vector pCAGGS containing the receptor binding domain (RBD, residues 319-541) of the spike glycoprotein from SARS-CoV-2, Wuhan-Hu-1 was obtained from BEI resources. Individual K417N, E484K, and N501Y substitutions were introduced to the coding sequenceby site-directed mutagenesis. The RBD variant containing three substitutions (K417N/E484K/N5101Y) was synthesized and cloned by Genscript into the pcDNA3.1 vector and subcloned into pCAGGS. Vectors were used to transform DH5α bacteria for transfection DNA prep. DNA was purified using the PureLink plasmid midiprep kit (Invitrogen, CA), filtered through a 0.22 μm filter, and stored at −20 °C. The SARS-CoV-2 spike glycoprotein RBD was expressed in Freestyle 293 cells (ThermoFisher, MA) grown in Freestyle 293 media at 37 °C with 8% CO_2_ and shaking at 135 rpm in a humidified incubator. Cells were transfected with plasmid DNA and 25 kDa polyethyleneimine (Polysciences, PA) at 1 μg/mL in Freestyle 293 media. Media was harvested 4 days after transfection and clarified by centrifugation at 4000 xg for 10 minutes. His-tagged RBD protein was purified from cell culture media by Ni-NTA chromatography and an AKTA pure system. Columns were washed with Buffer A (20 mM Sodium Phosphate pH 7.2, 500 mM NaCl) and bound protein was eluted with Buffer B (Buffer A + 250 mM Imidazole). Protein was then desalted using a HiPrep 26/10 desalting column and phosphate-buffered saline (PBS) pH 7.4 and concentrated using a 10 kDa centrifugal concentrator (Milipore-Sigma, MA). Protein concentration was determined by absorbance at 280 nm and an estimated extinction coefficient of 33850 M^−1^ cm^−1^. Concentrated protein was aliquoted, snap-frozen in liquid nitrogen, and stored at −80 °C.

### Unfolding Experiments

Guanine unfolding experiments were performed by fluorescent dye assay using bis-ANS (4,4’-dianilino-1,1’-binaphthyl-5,5’-disulfonic acid, Tocris, Bristol, UK) as described previously(21) with minor modifications. Protein (5 μM) was mixed with bis-ANS (10 μM) in PBS and guanidine HCl (prepared in PBS) ranging from 0 M to 2.5 M in a 96-well black plate in duplicate. Mixtures were incubated for 1 hour at room temperature before fluorescence readings were taken using a GloMax Explorer (Promega, WI) plate reader with an excitation filter at 405 nm and emission filter at 500 – 550 nm. Fluorescence data were used to calculate the fraction folded at a given guanidine concentration and analyzed by non-linear regression (22) to estimate the free energy of unfolding and best-fit values were compared by one-way ANOVA. Data presented are representative of three independent experiments using the same preparation of RBD. Temperature induced unfolding experiments were performed as described previously for ovalbumin(20). Briefly 5 μM RBD protein was mixed with 50 μM bis-ANS in 50 μL of 10 mM sodium phosphate buffer at pH 7.2 in a 200 μL PCR plate which was then sealed with a plate sealer. The plate was heated from 10 °C to 90 °C at a rate of 0.016 °C per second and bis-ANS fluorescence was recorded. Fluorescence data was plotted as a function of temperature and the peak was taken as the melting point. Tm values were compared by one-way ANOVA.

### RBD-ACE2 Binding Competition Assay

RBD-ACE2 binding competition assay was developed using the SARS-CoV-2 surrogate virus neutralization test kit (Genscript, NJ). First a 5-fold dilution series of RBD variant starting at 10 μM was prepared in sample dilution buffer in duplicate. Next, the serum/antibody incubation step was replaced by incubating horseradish peroxidase-labeled RBD (RBD-HRP) 1:1 with diluted RBD variant and incubated at 37 °C for 30 minutes. Then the kit procedure was followed as described in the manufacturer’s instructions. After development absorbance at 450 nm was measured with the GloMax Explorer plate reader. Absorbance values were plotted as a function of RBD variant concentration and analyzed by non-linear regression to estimate LogIC50 values. Best-fit values were compared by one-way ANOVA. Data presented are representative of three independent experiments.

### Limited Proteolysis

Limited proteolysis experiments were conducted with recombinant human cathepsin S (Milipore-Sigma) diluted in phosphate-citrate buffer at pH 5.6 and 37 °C(21). Proteolysis reactions were prepared in 150 μL phosphate-citrate buffer pH 5.6 containing 0.5 μg/μL of RBD variant, 0.025 μg/μL Cathepsin S, and 2 mM dithiothreitol. Reactions were prepared in triplicate and sampled after 0, 15, 30, and 60 minutes of incubation. Proteolysis was halted by mixing with sodium dodecyl sulfate-polyacrylamide gel electrophoresis (SDS-PAGE) loading buffer containing 150 mM 2-mercaptoethanol and incubation at 95 °C for 5 minutes. Proteolysis reactions were analyzed by SDS-PAGE and staining with Coomassie blue. Gels were imaged with a Chemidoc MP imaging system (Bio-Rad, CA) and analyzed with ImageJ software. Band intensities of the intact RBD were extracted from images and normalized to the 0-minute band intensity before analysis by two-way ANOVA.

## Acknowledgments

The following reagent was produced under HHSN272201400008C and obtained through BEI Resources, NIAID, NIH: Vector pCAGGS Containing the SARS-Related Coronavirus 2, Wuhan-Hu-1 Spike Glycoprotein Receptor Binding Domain (RBD), NR-52309.

---

**Figure S1.**
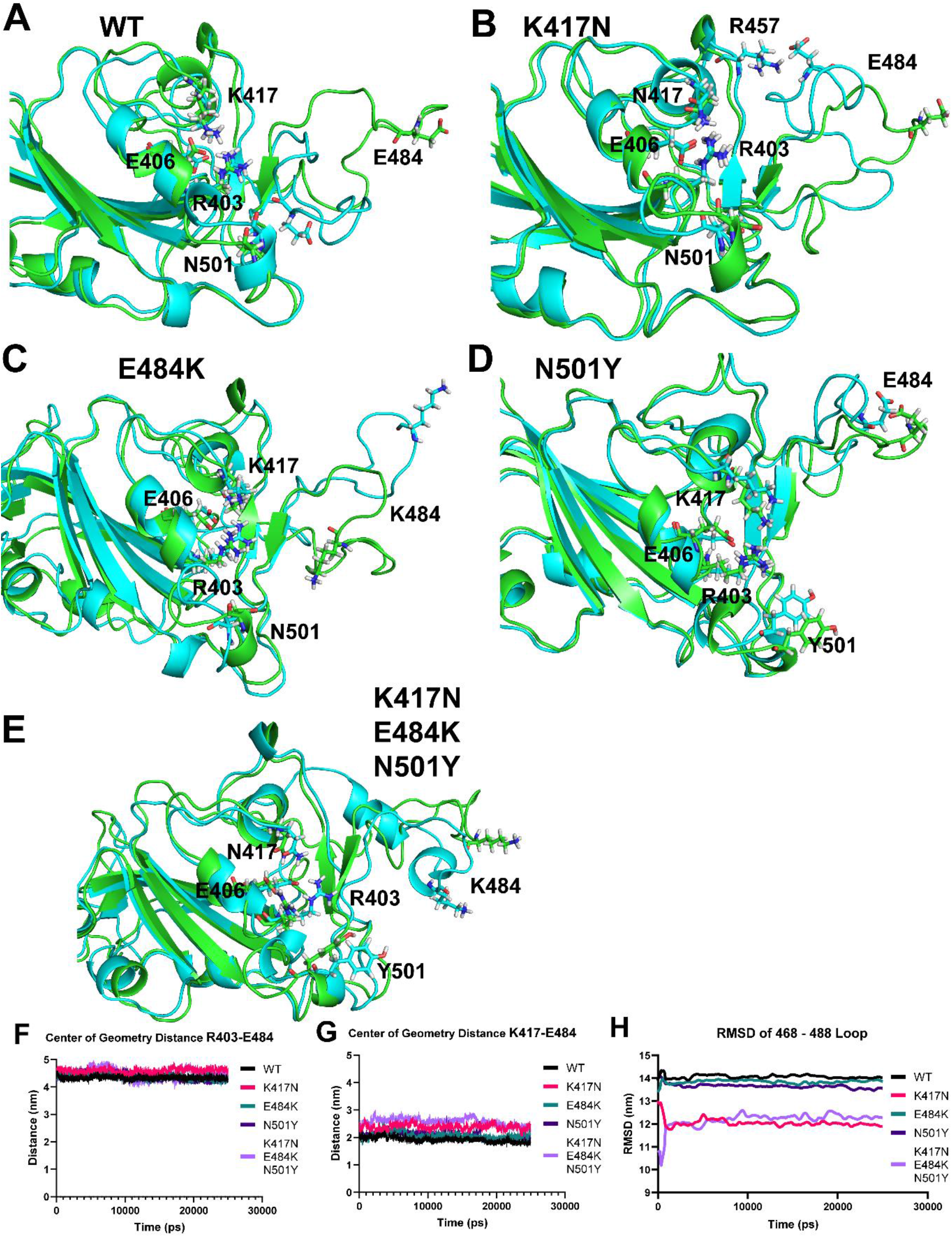
ff14SB specific changes in RBD residue contacts. A - E. RBD coordinates extracted from the molecular dynamics trajectories of the wild-type (A), K417N (B), E484K (C), N501Y (D) and K417N/E484K/N501Y (E) RBD variants in the ff14SB force field. The substituted residues as well as important hydrogen bonding pairs are shown. F. Center of geometry distance in nm between residue R403 and E484 plotted as a function of time for all RBD variants in the CHARMM36 force field. G. Center of geometry distance in nm between residue K417 and E484 plotted as a function of time for all RBD variants in the CHARMM36 force field. H. RMSD of the 468 – 488 loop relative to the starting structure for all RBD variants plotted as a function of time.

